# Genetic basis and adaptive implications of temperature-dependent and temperature-independent effects of drought on chickpea phenology

**DOI:** 10.1101/2022.01.26.477940

**Authors:** Yongle Li, Lachlan Lake, Yashvir S. Chauhan, Julian Taylor, Victor O. Sadras

**Affiliations:** School of Agriculture, Food and Wine, The University of Adelaide; South Australian Research and Development Institute, Australia; Department of Agriculture and Fisheries, Kingaroy, Australia

**Keywords:** carbon isotope, climate change, development, drought, flowering, genome, heat, phenotype, temperature, trade-off

## Abstract

Water deficit often hastens flowering of pulses partially because droughted plants are hotter. Separating temperature-independent and temperature-dependent effects of drought is important to understand, model and manipulate phenology genetically and agronomically.

We define a new trait, *drought effect on phenology (DEP =* difference in flowering time between irrigated and rainfed crops), and use F_ST_ genome scan to probe for genomic regions under selection for this trait. Genomic regions overlapping for early- and late-sown crops would associate with temperature-independent effects and non-overlapping genomic regions would associate with temperature-dependent effects.

Time to flowering shortened with increasing water stress quantified with carbon isotope composition. Genomic regions on chromosomes 4, 5, 7 and 8 were under selection for *DEP*. An overlapping region for early and late sowing on chromosome 8 revealed a temperature-independent effect with four candidate genes: *BAM1, BAM2, HSL2* and *ANT*. The non-overlapping regions included six candidate genes: *EMF1, EMF2*, BRC1/TCP18, *BZR1, NPGR1* and *ERF1*.

Modelling to assess *DEP* adaptive value showed it reduces the likelihood of drought and heat stress at the expense of cold risk. Accounting for *DEP* would improve phenology models to predict adaptation to future climates and breeding against the combined risks of drought, heat, and cold stress.

**Highlight:** Predictive and genetic models that overlook drought effects on phenology can return biased predictions of adaptation to future climates. Here we study the genetic causes and adaptive consequences of hastened flowering under drought.

## Introduction

Phenological shifts are the most conspicuous biological effects of global change, and the relative phenology of plants, herbivores and predators is central to the assemblage of trophic webs in natural and agricultural systems (Kankaanpää *et al*., 2020; Levine *et al*., 2002; Otegui *et al*., 2021; Parmesan, 2006; Richardson *et al*., 2013). Darwin (1859) observed “…very trifling changes, such as a little more or less water at some particular period of growth, will determine whether or not the plant sets a grain…”. This notion of a critical developmental period for seed production is central to plant physiology and agriculture, as farmers pair genotype and sowing time to manipulate crop phenology against the risks of frost, heat, drought, herbivory, and disease (Berger *et al*., 2006; Berger *et al*., 2004; Hunt *et al*., 2019; Lake *et al*., 2021; Otegui *et al*., 2021).

Temperature and photoperiod modulate the transition from the vegetative to reproductive stage and are at the core of predictive models (Lake *et al*., 2021; Mauney, 1963; Patrick and Stoddard, 2010; Summerfield *et al*., 1985; Wallach *et al*., 2021; Zheng *et al*., 2013). Fragmented empirical evidence shows that water deficit often hastens flowering in temperate grain legumes including chickpea (*Cicer arietinum, 2n = 2x = 16)*, the focus of this study (Anbazhagan *et al*., 2015; Fang *et al*., 2011; Johansen *et al*., 1994; Lizarazo *et al*., 2017; Singh, 1991; Thomas *et al*., 2004). Genotypic variation in this response is largely unexplored. Likewise, the adaptive and agronomic value of hastened flowering in response to water deficit is unknown but is expected to vary with soil and climate driving the patterns of supply and demand of water (Jordan and Miller, 1980; Schwinning and Ehleringer, 2001; Tardieu, 2012). Few *ad-hoc* models capture the effect of drought on flowering time (Chauhan *et al*., 2019; Lizarazo *et al*., 2017; McMaster *et al*., 2011) but mainstream crop models commonly used in climate change predictions do not (Wallach *et al*., 2021). Overlooking the effect of plant water status on phenology can therefore bias predictions of crop adaptation to future climates.

In contrast to the hastening of flowering in droughted chickpea (Chauhan *et al*., 2019; Fang *et al*., 2011), water deficit delayed time to first flower in Tunisian populations of burr medic (*Medicago polymorpha*) from wet (664 mm annual rainfall) and intermediate (345 mm yr^-1^) environments, with no effect on fast-developing populations from dry environments (173 mm yr^-1^) (Yousfi *et al*., 2015). The discrepancy between drought hastening or delaying development can be related to species, ecotype, and other factors such as the intensity of stress and interactions between water stress and temperature. For example, wheat (*Triticum aestivum*) phenological development responds non-linearly to plant water status, with mild water stress shortening and severe stress extending the time from floral initiation to anthesis (Angus and Moncur, 1977). In a factorial combining water regime and sowing time, water deficit hastened the flowering of mungbean (*Vigna radiata*) in early but not in late sowing, highlighting the interaction of water and temperature in modulating development (Thomas *et al*., 2004). Owing to the shift from latent heat to sensible heat, crop canopies are hotter under drought (Jones, 1992). Hence, hotter plant tissue may partially account for the effect of water deficit on phenology, but temperature-independent effects cannot be disregarded (McMaster *et al*., 2011). Separating temperature-dependent and temperature-independent effects of water deficit is important to understand, model and manipulate plant phenology genetically and agronomically.

Natural and agronomic selection may leave fingerprints in the genome, such as an extended genomic region where selection hitchhiking reduces diversity (Nielsen *et al*., 2005). The small genome of chickpea allows for whole-genome resequencing of contrasting genotypes to identify genomic regions under selection for agronomic traits (Li *et al*., 2017; Sadras *et al*., 2016). F_ST_ genome scan, where F_ST_ is the fixation index (Wright, 1950), uses a large number of molecular markers to scan regions with extreme genetic differentiation between diverging populations (Fumagalli *et al*., 2013; Holsinger and Weir, 2009). F_ST_ genome-scan is based on neutral theory, assuming that polymorphisms are selectively neutral and random genetic drift is the main driver of allele frequencies in populations without selection (Booker *et al*., 2020). This approach to detect selection signals in small samples has been insightfull in ecological and evolutionary settings (Barr *et al*., 2021; Van Bocxlaer, 2017), for crops including rice (*Oryza sativa*), wheat and chickpea (Jordan *et al*., 2015; Li *et al*., 2017; Sadras *et al*., 2016; Xu *et al*., 2012), and for crop pests such as the soybean aphid, *Aphis glycines* (Coates *et al*., 2020). The reliability of F_ST_ genome scan is particularly apparent in a comparison between F_ST_ genome scan and genome-wide association (GWAS), with both returning a common 100 kb region (AB4.1) on chromosome 4 associated with Ascochyta blight resistance in chickpea (Li *et al*., 2017).

Here we define a new trait, *drought effect on phenology (DEP =* difference in flowering time between irrigated and rainfed crops), to test three hypotheses in a study combining field experiments, F_ST_ genome scan, and modelling. First, time to flowering is shortened in proportion to plant water deficit, and this response is genotye-dependent. Second, the effect of drought on phenology involves genes associated with both temperature-independent and temperature-dependent components. Genomic regions under selction for *DEP* that are common to early- and late-sown crops would support temperature-independent effects while non-common genomic regions would indicate temperature-dependent effects. Third, drought modulation of phenology drives a site-dependent reduction in drought and heat stress at the expense of cold stress; this hypothesis was tested with modelling in a north-south transect with varying soils, rainfall and thermal regimes in eastern Australia.

## Methods

### Phenotyping phenology, carbon isotope composition and seed size in the field

A field experiment was established on a calcic luvisol (Isbell, 1996) at Roseworthy, South Australia (34°52’ S, 138°69’E) that combined factorially 20 genotypes (Table 1), two water regimes (dry, rainfed; wet, sprinkler irrigated), and two sowing times (early, early June; late, early-mid July). The experiment was repeated twice over successive seasons. Treatments were laid out in a split-split-plot design with three replicates, where sowing time was assigned to the main plot, water regime to the sub-plot, and genotypes randomised within each plot. Each plot comprised 6 rows, 0.24 m appart, 5-m long. Further details of the experiment are in Sadras et al. (2016).

**Table 1.**
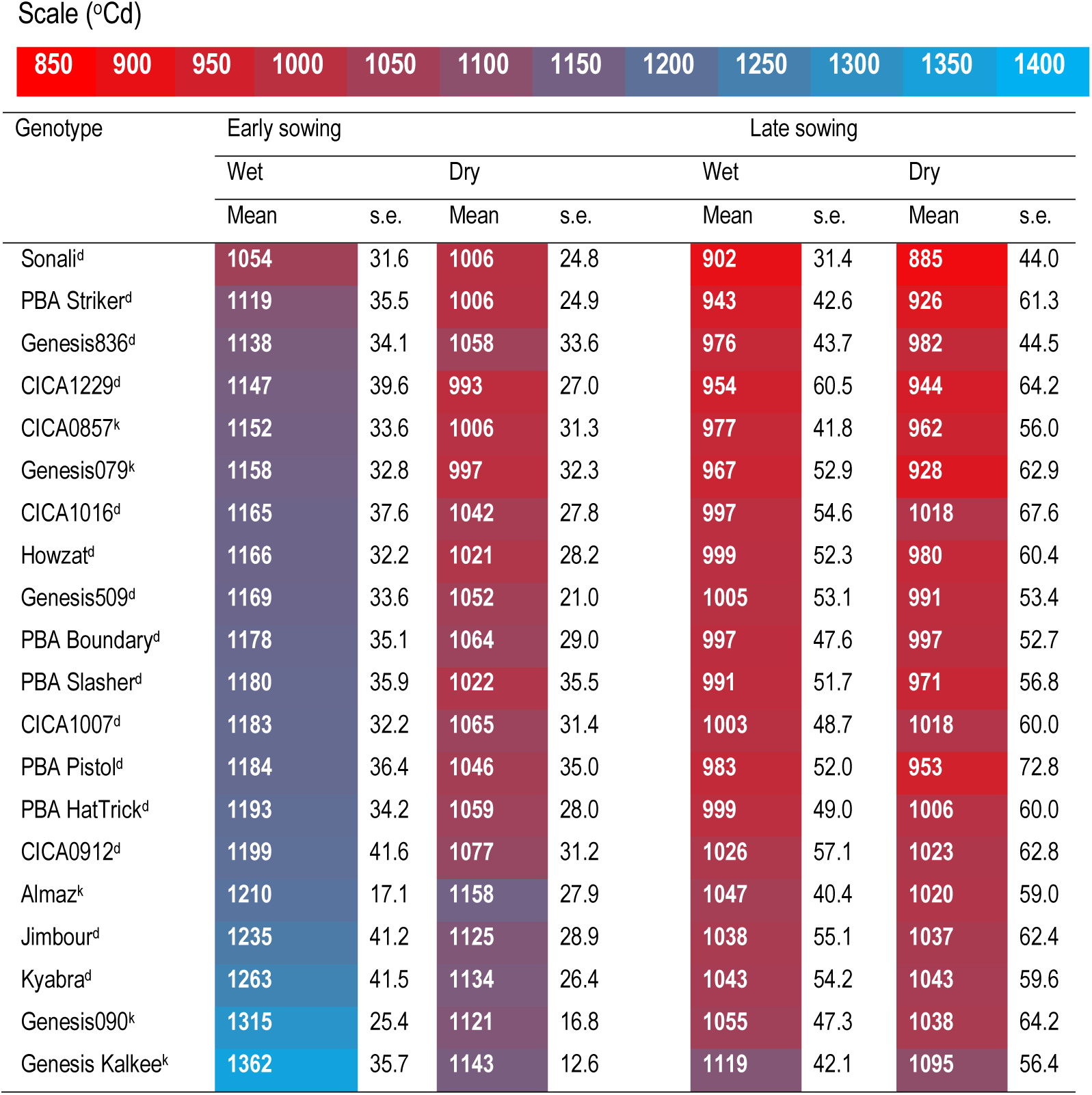
Thermal time (°Cd) from sowing to flowering in 20 chickpea genotypes grown under two water regimes (wet, dry) and two sowing times (early, late). Superscripts indicate (d) Desi and (k) Kabuli genotypes. Data are averaged for two seasons.

To avoid bias associated with border effects, all measurements were made in the center rows (Rebetzke *et al*., 2014). We scored phenology weekly to establish the time to 50% of plants at beginning of flowering and calculated thermal time from sowing to flowering using a base temperature of 0 °C (Berger *et al*., 2006; Berger *et al*., 2004). To quantify crop water status, we measued carbon isotope composition (δ^13^C) at peak biomass, shortly after flowering. This trait integrates crop water status over the growing period until sampling time and is robust in relation to environmental conditions – radiation, wind speed, temperature, vapour pressure deficit (Condon *et al*., 2002), unlike traits such as stomatal conductance, leaf water potential or canopy temperature that vary with conditions at sampling time. Ten shoots per replicate were sampled and dried at 70 °C for 48 h; subsamples were ground and analysed for C isotope composition using a Europa 20-20 stable isotope ratio mass spectrometer with an ANCA-SL (Automated Nitrogen Carbon Analysis for Solids and Liquids) preparation system. In a batch of samples, after every eighth sample a test and a reference (Pee Dee Belemnite) were determined and used to correct for any drift or carryover in the instrument. Carbon isotope composition δ^13^C was calculated as (Condon et al., 2002):

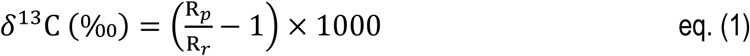

where R is the ^13^C/^12^C ratio and subscripts indicate plant (p) and reference (r).

To probe for associations between phenology and seed weight, as expected from pleiotropic effects (Hovav *et al*., 2003; Kumar and Abbo, 2001), we measured average seed weight at maturity after drying and threshing 2-m^2^ plant samples.

### Statistical analysis of crop traits

Time from sowing to flowering, carbon isotope composition and seed weight were analysed with a linear mixed model (LMM) where the fixed component consisted of the crossed factor combinations of genotype, sowing time, water regime and season. This ensured the LMM fixed effects included main effects for each of the factors as well as the full complement of second, third and fourth order interactions. Additional sources of variation associated with aspects of the field design such as replicates, or non-linear trends across the rows or ranges of the experiment, were modelled using random effects. Due to the distinct sowing times within each season, and the potential for traits to vary significantly between water regimes, we partitioned the model residuals of the LMM to ensure a separate residual variance was specified for each combination of season by sowing time by water regime. From this complete model, Wald ANOVA tables were extracted for summary. To appropriately compare genotypes between water regimes, BLUEs for genotype by water regime by sowing time, and averaged over season, were predicted from the LMM.

Generalised broad-sense heritability was calculated for all traits using the method by Cullis et al. (2006). This involved re-fitting the LMM with the genotype factor as a random effect and leaving other terms unchanged in the LMM specification. Heritabilities are then considered to be averaged over water regimes, sowing times and seasons.

To explore associations between variables, we fitted least square regression (Model I) when error in *x* was negligible in comparison to error in *y* and reduced maximum axis regression (RMA, Model II) to account for error in both *x* and *y* (Niklas, 1994). For both ANOVA and regressions, we present *p* as continuous values, avoiding arbitrary *p* thresholds for significance (Wasserstein *et al*., 2019).

*Drought effect on phenology (DEP)* was calculated in two ways, with difference- and residual-based approaches. First, we calculated *DEP* as the *difference* in flowering time between the dry and wet treatments. A reduced LMM was fitted using the above model, with the water regime treatment omitted. From this LMM, BLUEs for the genotypes within each sowing time were predicted. The second approach uses the BLUEs of flowering time by genotype, sowing time and water regime extracted from the full fitted LMM defined above. Within each sowing time, BLUEs of flowering time for the wet treatment were regressed against the BLUEs of flowering time for the dry treatment, and the *residuals* from the RMA regressions were taken as a proxy for *DEP* (Erena *et al*., 2021; McDonald *et al*., 2018). *DEP* calculated as differences correlated closely with *DEP* calculated as residuals (r = 0.93 for first sowing, r = 0.96 for second sowing, p < 0.0001 for both; Supplementary Figure 1). Hereafter, we report difference-based *DEP* for clearer biological interpretation; for example, early sown Genesis Kalkee returned a difference-based *DEP* = 219 °Cd, which means drought hastened flowering by 219 °Cd in relation to well-watered crops.

### DNA sequencing and *F_ST_* genome scan

DNA extraction and sequencing have been described previously (Sadras *et al*., 2016). Briefly, we extracted DNA of the 20 chickpea genotypes from young leaves using Qiagen DNeasy Plant Mini Kit. TruSeq libaries were constructed for each genotype with an insert size of 500 base pairs and sequenced using Illumina HiSeq 2000 platform. Pair-end reads (100 bp) were trimmed and mapped to the reference genome 2.6.3 (http://cicer.info) using SOAP2 (Li et al., 2009). To perform F_ST_ genome scan, the BAM files of the top six and bottom six genotypes based on the adjusted entry means of *DEP* were selected as contrasting populations for F_ST_ estimation. F_ST_ of the two contrasting populations were estimated using software ngsTools and ANGSD (Fumagalli *et al*., 2013; Fumagalli *et al*., 2014; Korneliussen *et al*., 2014). F_ST_ is a measurement of genetic differentiation between populations, with larger F_ST_ indicative of larger divergence between the populations. The whole genome was scanned for each 100 kb window (non-overlapping) to find regions with extreme F_ST_ (compared with the adjacent regions) as an indicator of regions under selection. The assumption is that if a region is under selection, the pattern of genetic differentiation between populations may change, i.e. alleles may be fixed in a particular population. Genomic regions with the top 0.1% F_ST_ were considered to be under selection (Sadras *et al*., 2016).

### Modelling the adaptive value of drought effect on phenology

Current models reliably predict phenology but not yield of pulses particularly because algorithms are lacking that relate yield and extreme temperatures (Lake *et al*., 2021). Thus, to test our third hypothesis, we modelled the phenology of two contrasting genotypes – responsive vs. unresponsive to water deficit – to quantify the phenology-driven differences in water stress and temperature in the critical period from flowering to 200 °Cd after flowering (Lake and Sadras, 2014). The expectation is reduced water stress and lower temperature during the critical period of the responsive genotype in relation to its unresponsive counterpart (Fig. 1a). We used APSIM (Classic version 7.10) to simulate flowering time and the daily water stress index, *WSI*. The *WSI* is the ratio between the potential water supply, which depends on the volume and wetness of soil explored by roots, and the water demand of the canopy, which is a function of radiation, ambient temperature and humidity (Chenu *et al*., 2011).

**Figure 1.**
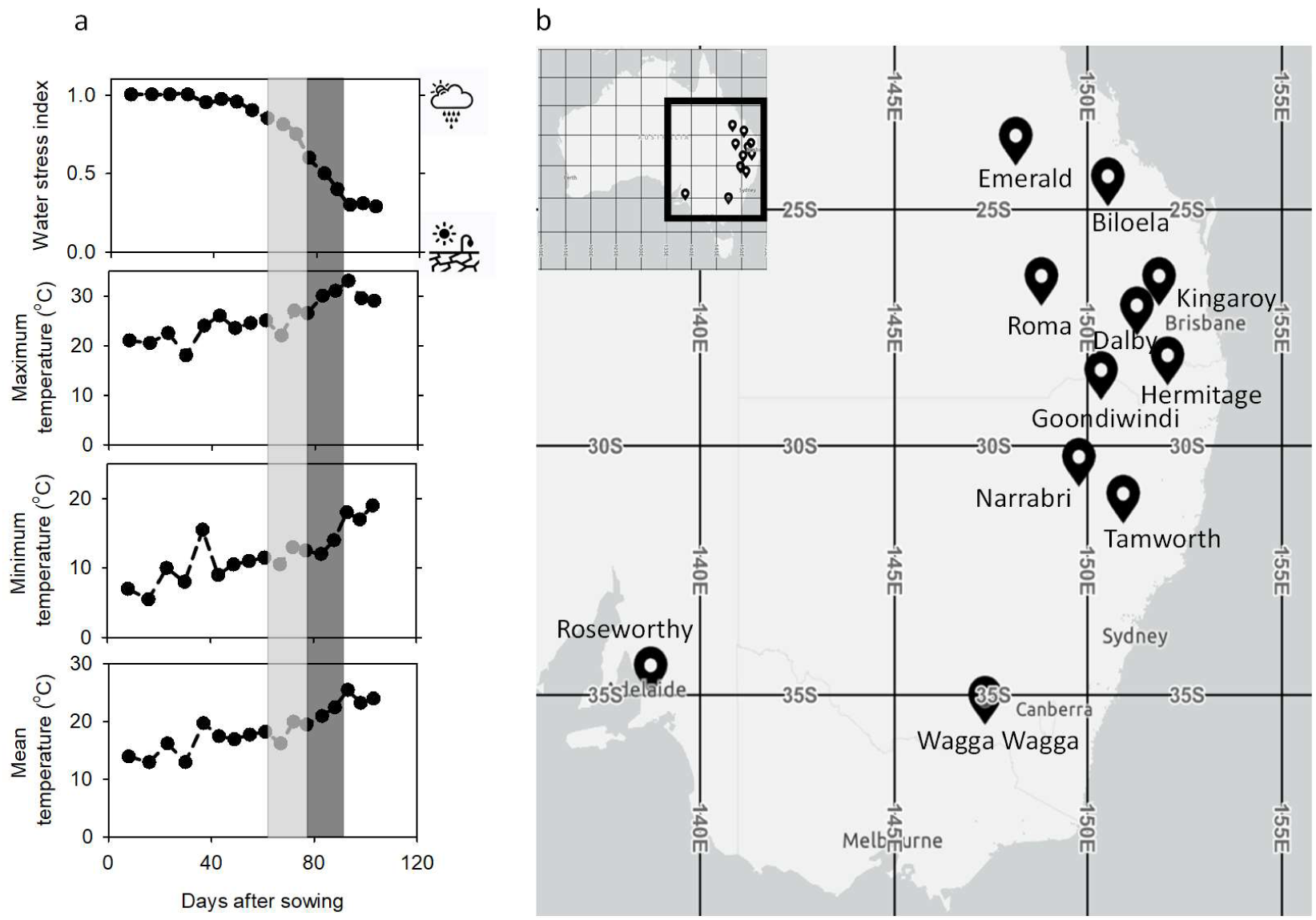
(a) Illustration of the dynamics of water stress index, maximum, minimum and mean temperature during the growing season of chickpea in relation to the critical period for a genotype responsive to water deficit (light grey) and an unresponsive genotype (dark grey). The water stress index ranges from 1 (no stress) to 0 (no growth). (b) Transect of locations in eastern Australia, from Emerald to Wagga Wagga, used to model the phenology, water stress index and temperature during the critical period; Roseworthy, the experimental site, was also included in the simulations.

The *WSI* ranges from 1 (no stress) to 0 (no growth) (Fig. 1a). APSIM is a widely used crop simulaton framework that has been extensively validated for multiple crops and environments in Australia and elsewhere (Holzworth *et al*., 2014; Keating *et al*., 2003). Tests of the model’s ability to simulate phenology and *WSI* are particularly relevant for our study. Modelled chickpea flowering time as a function of temperature, daylength and soil water content correlated closely with measured flowering time in eastern Australia (Chauhan *et al*., 2019). The *WSI* has been extensively used for spatial characterisation of drought in many crop species and environments (Chenu, 2015). For chickpea in Australia, the modelled *WSI* is biologically and agronomically robust as it defines drought types that correlate with seed yield (Chauhan *et al*., 2017; Lake *et al*., 2016).

We modelled two “isolines”, responsive and non-responsive to drought, using the same genetic parameters (Supplementary Table 1) except for phenological development of the responsive genotype for which developmental time was scaled with the algorighms developed and tested by Chauhan et al. (2019) to capture the drought effect on phenology. The two genotypes were compared in a factorial combining 11 locations in eastern Australia (Figure 1b), 65 years from 1957, five sowing times (at fortnigtly intervals from 14^th^ of May to 14^th^ of July) and two initial soil water contents (reset to field capacity or 50% field capacity on the 1^st^ of December of each preceding year). At sowing, a 20 mm irrigation was applied to ensure establishment. Climate data were sourced from Queensladn Goverment data base^1^. Phothethermal and rainfall regimes of these environments have been described in detail (Chauhan *et al*., 2017; Chauhan *et al*., 2008; Rodriguez and Sadras, 2007; Sadras and Rodriguez, 2007). Soil properties were obtained from the APSoil database (www.apsim.info). Out of the 7150 combinations in this factorial, 109 were failed crops as defined in Chauhan et al. (2017); the analysis thus focused on 7041 combinations. Using these data, the responsive and unresponsive genotypes were compared in two analyses. First, we calculated frequency distributions of flowering time, and *WSI*, maximum, minimum and mean temperature in the critical period. Second, for *WSI* and temepratures, one-to-one scatterplots were drawn against the null hypothesis of no difference between genotypes represented by the *y = x* line; deviations from *y = x* line were analysed with ANOVA to account for the effect of location, time of sowing and climate change. To assess the variation in the adaptive value of *DEP* with climate change, we used the updated World Meteorological Organisation climatological standard (Hulme, 2020); data were partitioned in ‘present-day’ climate, from 1991, and ‘historic’ climate before 1991.

## Results

### Photothermal and water regimes caused large variation in crop water status and phenology

Figure 2 summarises photothermal and water regimes. Solar radiation increased from 10.3-10.6 MJ m^-2^ for early sowing to 12.2-13.6 MJ m^-2^ with late sowing, maximum temperature from 16.8 to 18.6-19.9 °C and vapour pressure deficit from 0.70-0.73 kPa to 0.81-0.96 kPa (Fig. 1a).

**Figure 2.**
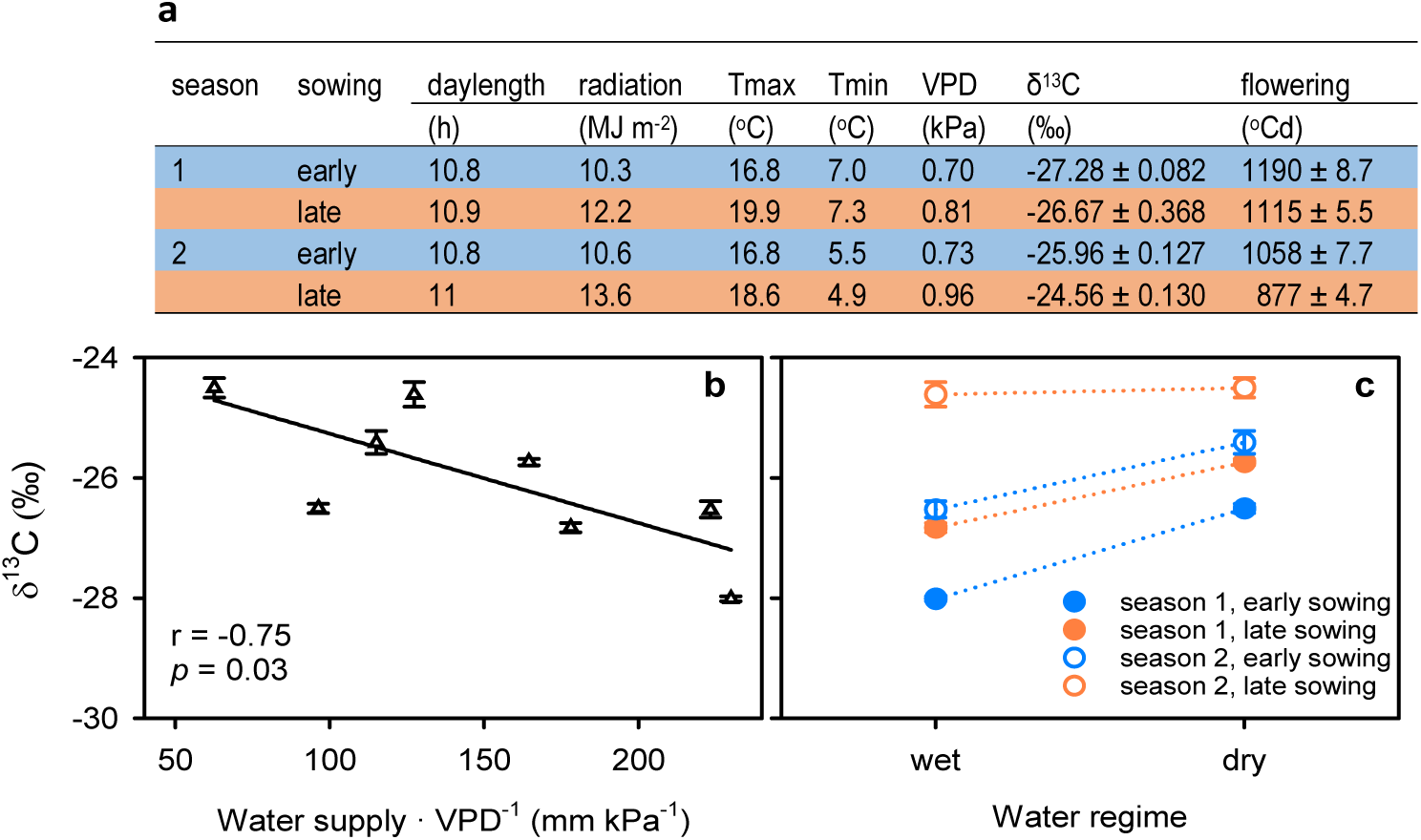
Photothermal and water regimes for chickpea crops associated with experimental sources of variation. (a) Daylength, solar radiation, maximum temperature (Tmax), minimum temperature (Tmin), vapour pressure deficit (VPD). Daylength is at sowing and the other weather variables are averages from sowing to average time of flowering of 20 genotypes. Carbon isotope composition (*δ*^*13*^*C)* and thermal time from sowing to flowering are averages (± s.e.) across 20 genotypes. (b) Relationship between carbon isotope composition (*δ*^*13*^*C)* and water supply : vapour pressure deficit ratio. Water supply and VPD are average from sowing to average time of flowering of 20 genotypes; owing to the lack of reliable measurement of plant available water at sowing, water supply was calculated as the sum of rainfall and irrigation. The line is the least square regression. (c) Variation in *δ*^*13*^*C* with water regime, season and sowing date. In b and c, *δ*^*13*^*C* is the environmental mean, calculated as the average of 20 genotypes. In b and c, error bars are two standard errors of the mean and are not shown when smaller than symbol.

Carbon isotope composition correlated with the ratio of water supply : demand (Fig. 2b). Carbon isotope composition indicated more severe water deficit in the dry than in the wet treatment and in late compared to early sowing (Fig. 2c, Supplementary Table 2). The difference in δ^13^C between wet and dry regimes was slightly smaller in late than in early-sown crops (Fig. 2c; Supplementary Table 2: water regime x sowing time interaction, *p* = 0.06).

Daylength varied from 10.8 to 11.0 h. (Fig. 2a). Detailed studies indicate this small variation in daylength can be regarded as a minor influence on phenology (Daba *et al*., 2016b; Hovav *et al*., 2003). Across genotypes, time from sowing to flowering ranged from 69 to 108 d or 877 to 1190 °Cd. Thermal time to flowering was unrelated to daylenght, and aligned with δ^13^C (r = −0.99, p = 0.003). Hence, variation in phenology with sowing time was mostly attributable to temperature and water regime, as discussed below.

### Hypothesis 1: time to flowering is shortened in proportion to plant water deficit and this effect is genotype-dependent

Broad-sense heritability of thermal time to flowering was 0.98; it varied with genotype, water regime and sowing time from 885 °Cd or 69 d for late-sown Sonali in the dry treatment to 1362 °Cd or 109 d for early-sown Genesis Kalkee in the wet treatment (Table 1, Supplementary Table 2).

Broad-sense heritability of δ^13^C was 0.78 and thermal time from sowing to flowering correlated strongly with δ^13^C (Figure 3a). A non-linear model improved slightly the correlation between thermal to flowering and δ^13^C, with p = 0.003 for the quadratic term (dashed line in Fig. 3a).

**Figure 3.**
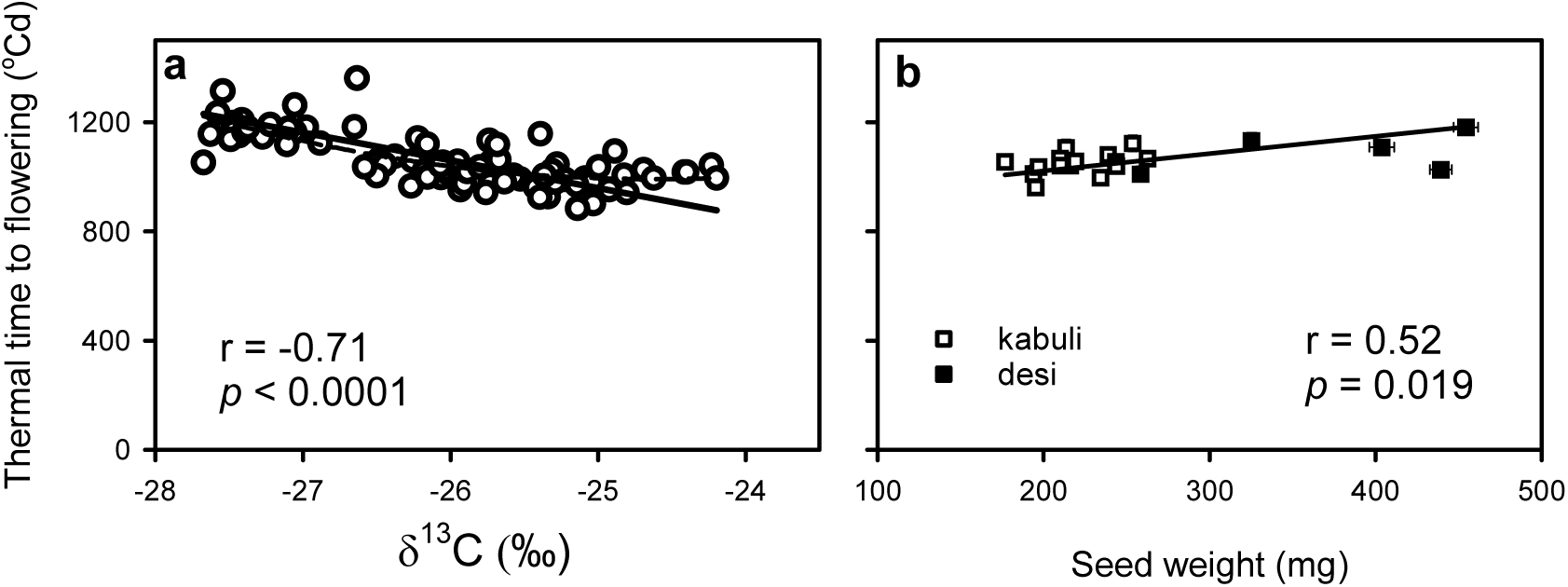
(a) Association between thermal time from sowing to 50% flowering and carbon isotope composition δ^13^C in 20 chickpea genotypes grown under two water regimes and two sowing dates across two seasons. (b) Association between thermal time from sowing to 50% flowering and average seed weight for 20 genotypes averaged across sources of variation. In b, error bars are two standard errors and are not shown when smaller than symbol. In a, b, the solid lines are reduced maximum axis regression (RMA, Model II) to account for error in both *x* and *y* (Niklas, 1994). The dashed line in (a) is a quadratic model with slightly higher r^2^ than the linear model (0.54 vs 0.49).

Broad-sense heritability of seed weight was 0.99. As expected from pleiotropic effects, thermal time to flowering correlated with seed weight; the slope of the RMA regression, 0.63 °Cd mg^-1^, could be useful for modelling (Fig. 3b).

Broad-sense heritability of drought effect on phenology was 0.61. This trait varied with genotype, sowing time, and with the interaction between genotype and sowing time: 4.6-fold in early-sown crops, from 47 °Cd in Sonali to 218 °Cd in Genesis Kalkee, and smaller (1.9-fold) variation in their late-sown counterparts (Fig. 4, Supplementary Table 3).

**Figure 4.**
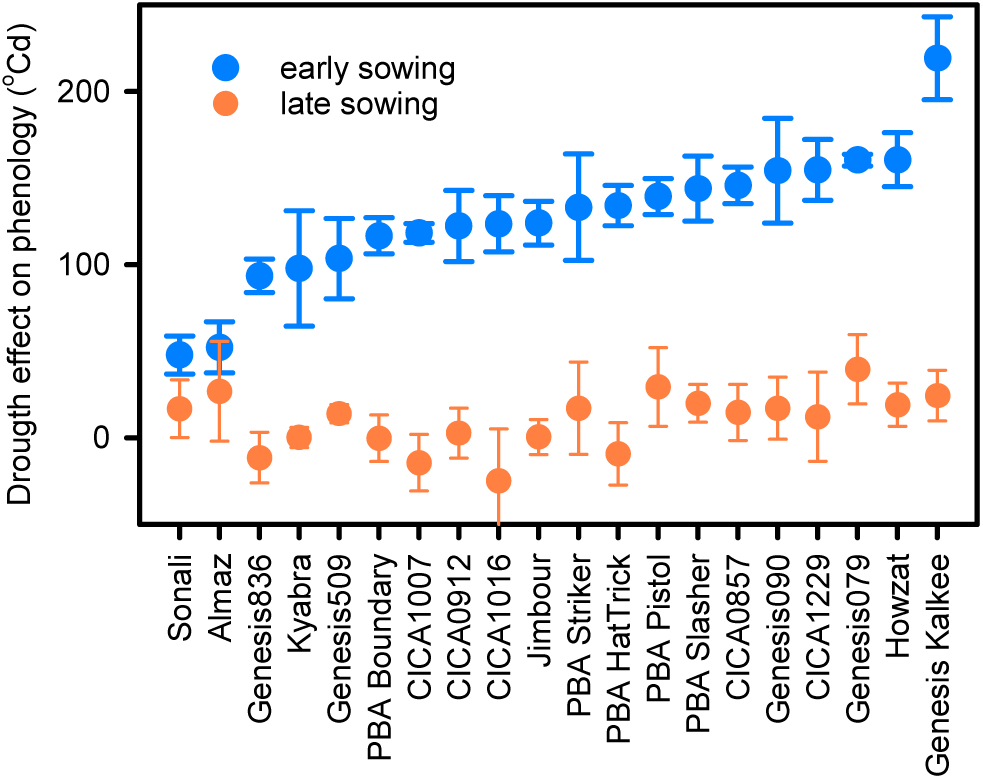
Genotype-dependent variation in drought effect on phenology, defined as the difference in thermal time from sowing to flowering between dry and wet treatments. Data are averaged across two experimental seasons and error bars are two standard errors.

### Hypothesis 2: *F_ST_* genome scan revealed genes associated with temperature-independent and temperature-dependent effects of drought on phenology

Figure 5 shows the F_ST_ genome scan to probe for the effects of drought on phenology, Supplementary Table 4 lists the genes located within 250kb of the genomic regions under selection (top 0.1% F_ST_) for this trait, and Table 2 summarises selected candidate genes. Genomic regions on chromosomes 4, 5, 7 and 8 were under selection for *DEP*. For early sowing, F_ST_ scan identified four genomic regions on chromosomes 5 and 8 that were under selection. For late sowing, four regions on chromosomes 4, 5, 6 and 8 were found under selection. Regions on chromosome 8 common to the two sowing times, which are only ∼100kb apart with high linkage disequilibrium among SNPs (r^2^ = 0.42), indicate a common set of genes associated with drought effect on phenology independent of temperature.

**Table 2.**
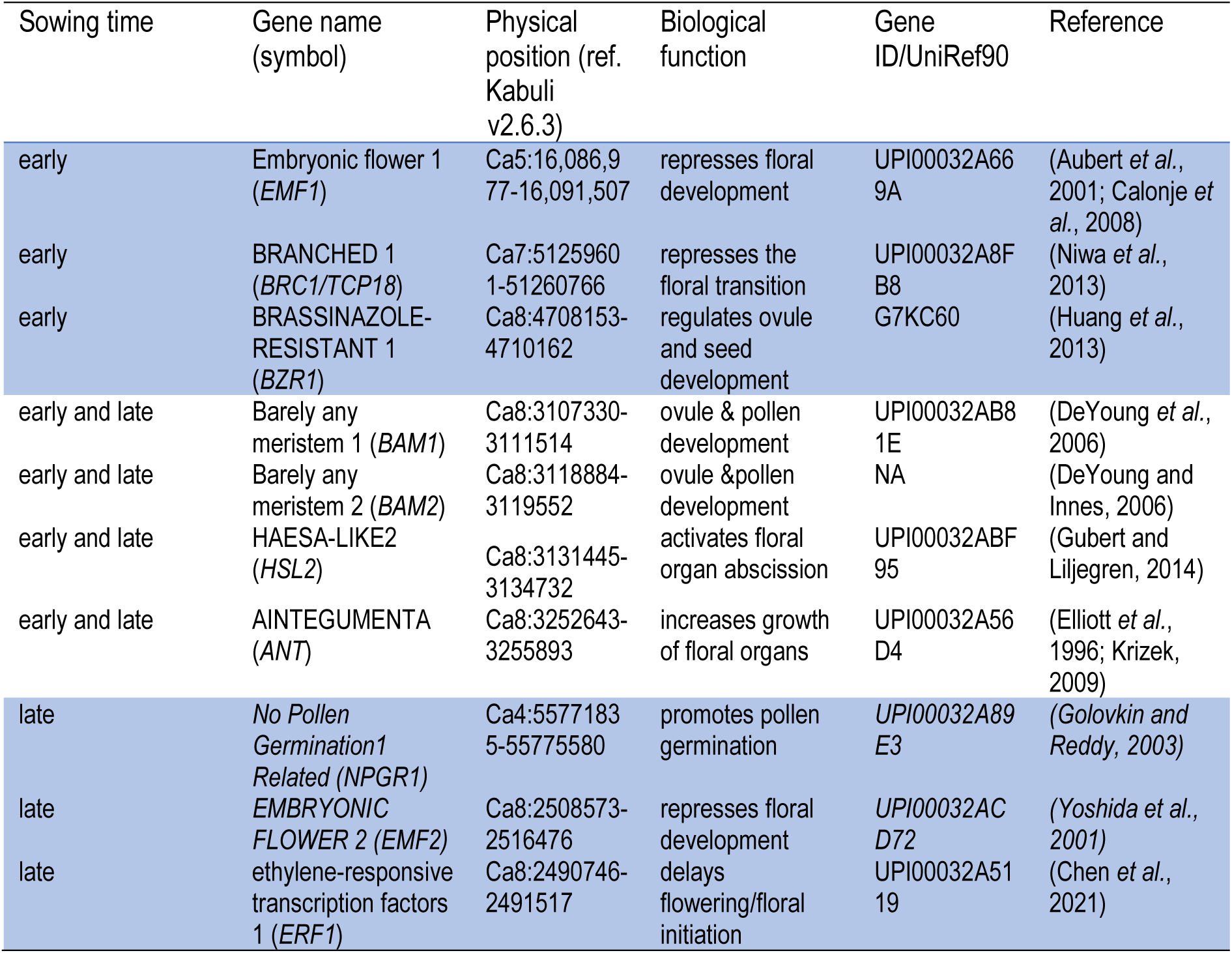
Selected candidate genes idenfitied to be under selection for drought effect on phenology. Three groups are considered: genes only identified in early- or late-sown crops (blue background), and those common to early- and late-sown crops.

**Figure 5.**
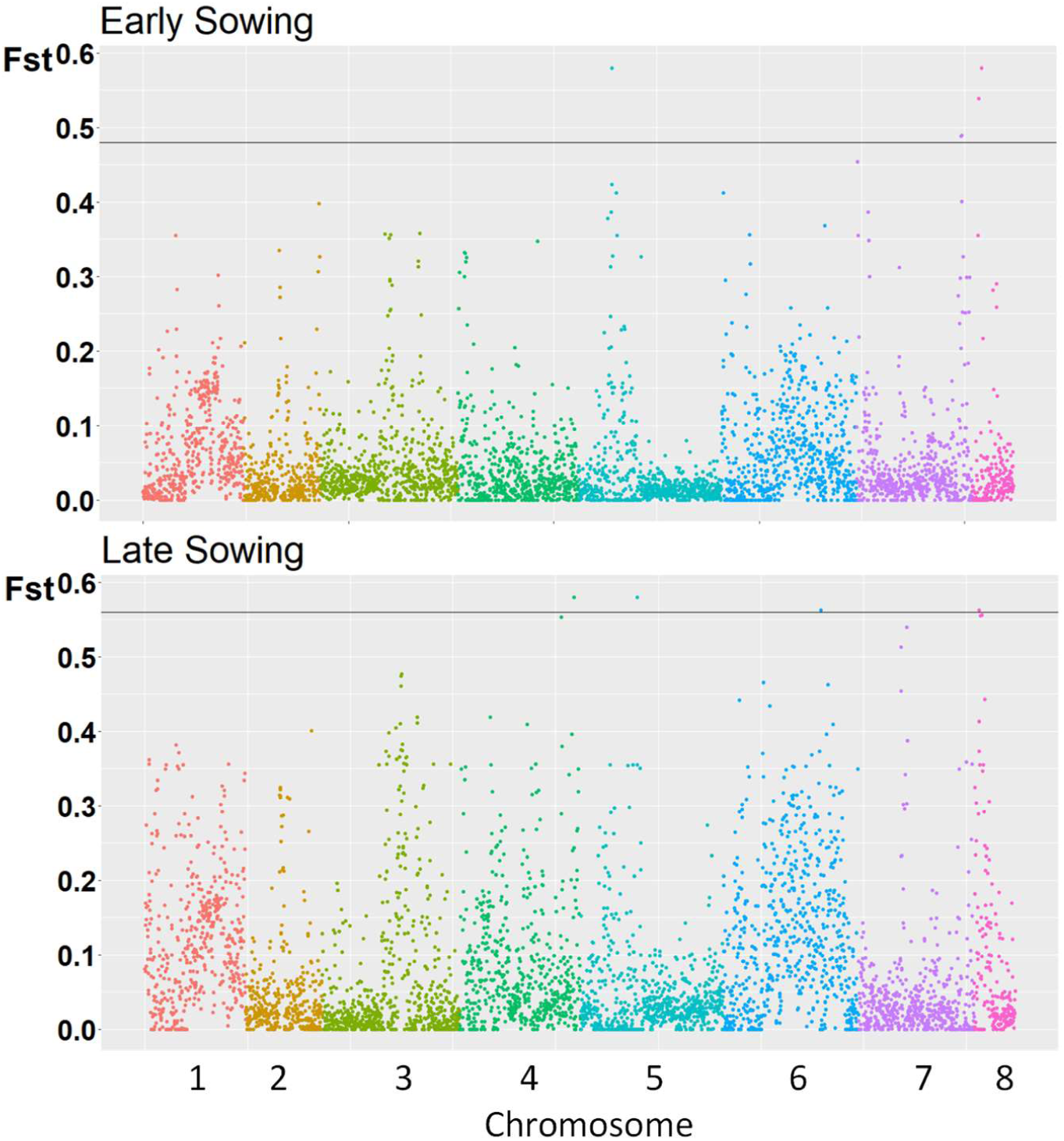
F_ST_ genome scan for drought effect on phenology in early and late sown chickpea. The x-axis corresponds to the eight chickpea chromosomes and each dot represents a F_ST_ estimated using SNPs from a window of 100 kb region. The horizontal line is the top 0.1% threshold. Dots above the threshold are genomic regions considered to be under selection for drought effect on phenology.

### Hypothesis 3. Water deficit modulation of phenology drives a site-dependent reduction in water stress and heat stress at the expense of cold risk in eastern Australia

Figure 6 shows histograms of modelled flowering time, water stress index and temperature in the critical period, and Supplementary Table 5 summarises the associated statistics. On average, time to flowering was 16 d or 227 °Cd shorter in the responsive genotype compared to its unresponsive counterpart.

**Figure 6.**
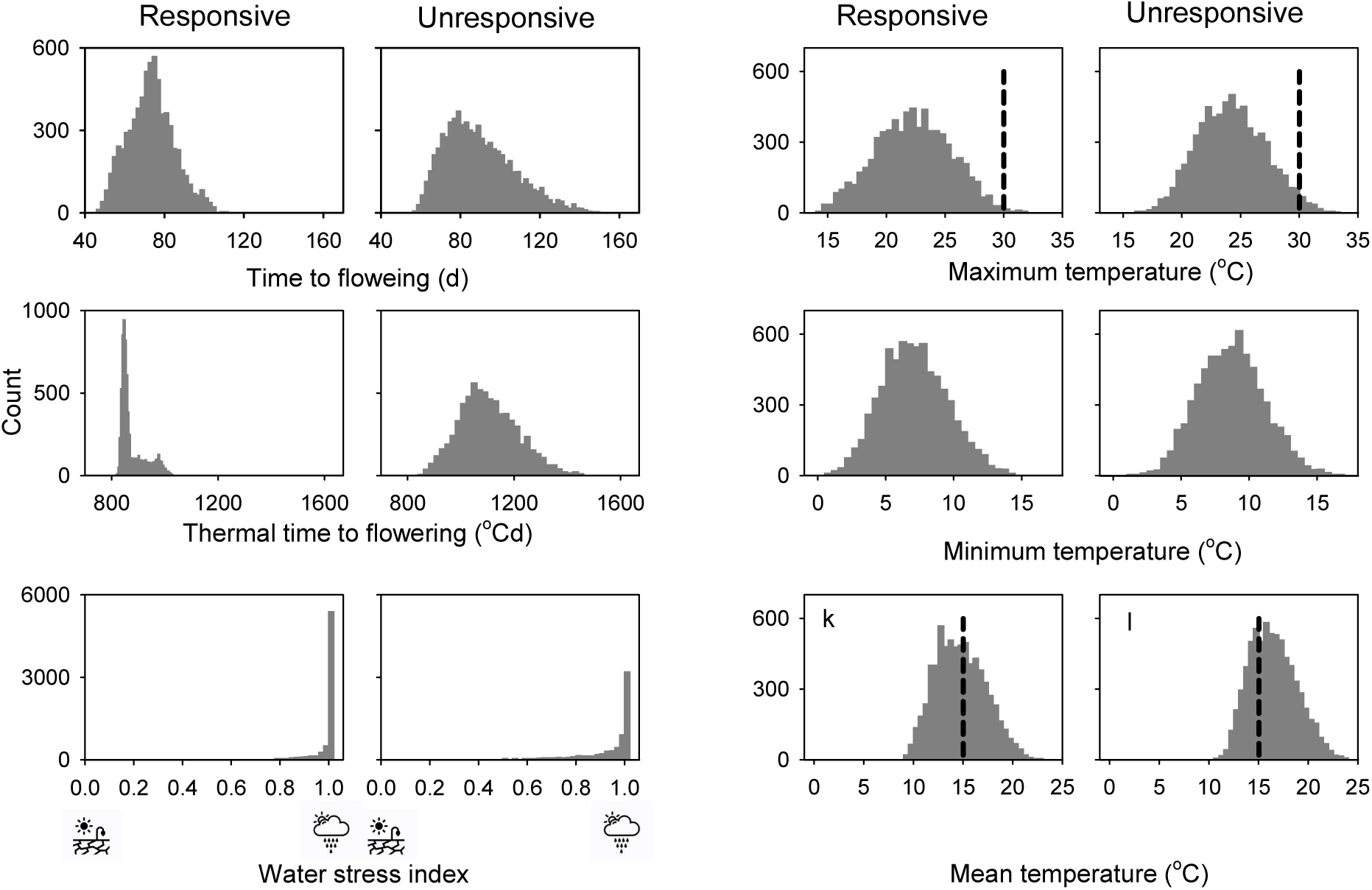
Frequency distribution of time from sowing to flowering and thermal time from sowing to flowering, and water stress index, maximum, minimum and mean temperature in the critical period of chickpea for a genotype phenologically responsive to water deficit and an unresponsive genotype. Modelled data from the combination of 11 locations, 65 seasons, two levels of initial soil water and five sowing dates (n = 7041, resulting from 7150 combinations minus 109 failed crops). The dashed lines are thresholds for reproductive disruption: above 30 °C for maximum temperature (Devasirvatham *et al*., 2012) and below 15 °C for mean temperature (Berger et al., 2012).

The frequency distribution of water stress index was J-shaped, with negative skewness and positive kurtosis. The number of cases with *WSI* = 1 (no stress) decreased from 5384 out of 7041 (76%) in the responsive genotype to 3186 (45%) for the unresponsive one.

Owing to the hastened phenology with water deficit, the responsive genotype averaged 1.9 °C lower maximum temperature, 1.6 °C lower minimum temperature and 1.7 °C lower mean temperature in the critical period than the unresponsive genotype. Maximum temperature over 30 °C (Devasirvatham *et al*., 2012) and mean temperature below 15 °C (Berger *et al*., 2012) in the critical period compromise chickpea reproduction. Maximum temperature over 30 °C was reduced from 183 cases in the unresponsive genotype to no cases for its responsive counterpart. Mean temperature below 15 °C increased from 2148 cases (31%) in the unresponsive genotype to 3073 cases (44%) for its responsive counterpart.

One-to-one comparisons help visualise the difference in water stress and temperature in the critical period between responsive and unresponsive genotypes (Fig. 7aei). Data close to the *y = x* line would indicate no difference between genotypes. The water stress index was on or below the *y = x* line in Fig. 7a, highlighting the consistent alleviation of drought associated with the hastening of flowering in the genotype responsive to water deficit. The one-to-one comparison showed most data were above the *y = x* line for both maximum and minimum temperature, reflecting the cooler critical period associated with earlier flowering in the responsive genotype (Fig. 7e, i). However, there were departures from this trend as the responsive genotype experienced lower maximum temperature than the unresponsive genotype in 839 cases (12%) and lower minimum temperature in 1170 cases (17%).

**Figure 7.**
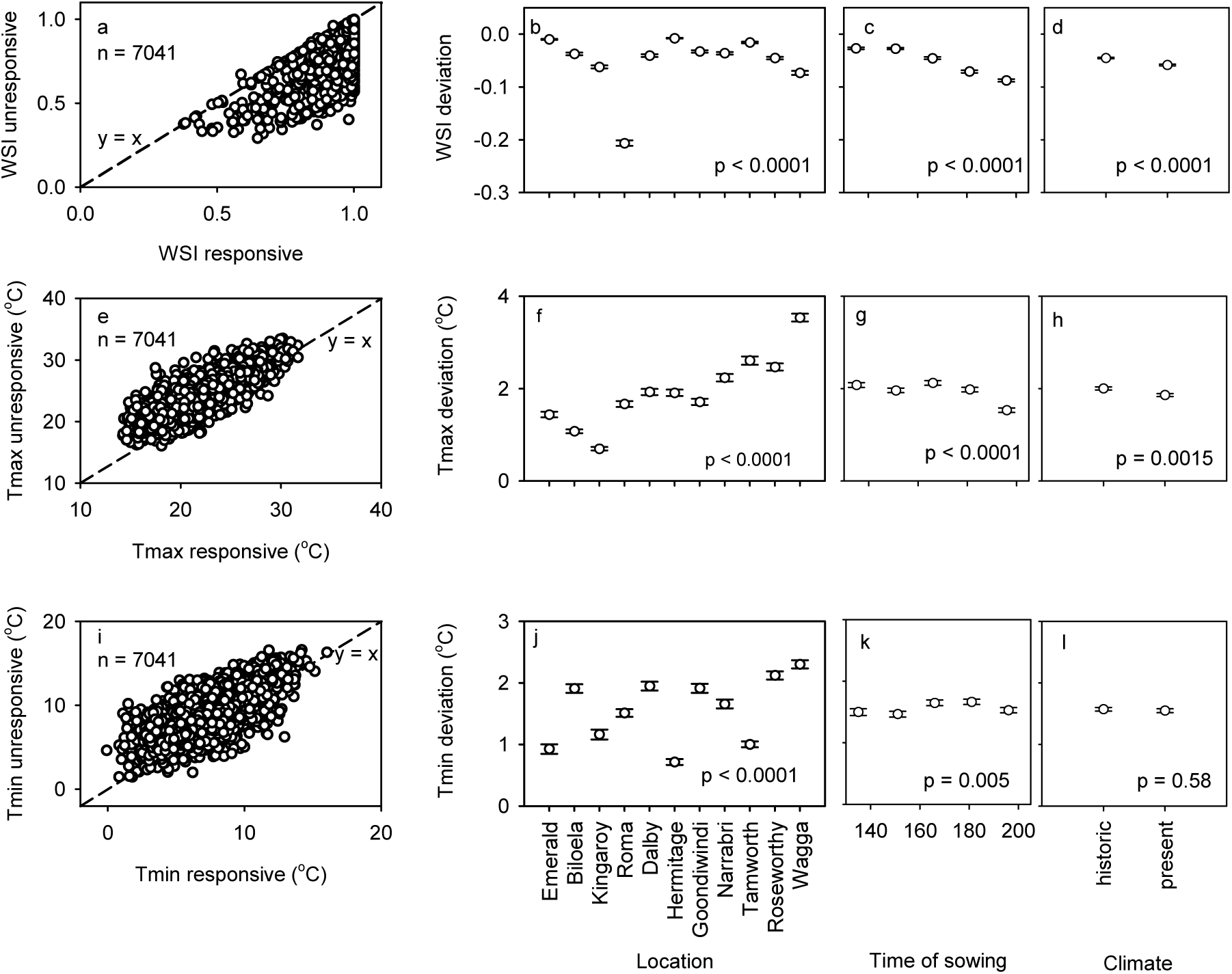
One-to-one comparisons of (a) water stress index, (e) maximum temperature and (i) minimum temperature during the critical period of chickpea between unresponsive and responsive genotypes. Modelled data are from the combination of 11 locations, 65 seasons, two levels of initial soil water and five sowing dates (n = 7041, resulting from 7150 combinations minus 109 failed crops). The dashed line, *y = x*, represents the null hypothesis of no difference between genotypes. Average deviation of *WSI* from the *y = x* line for (f) 11 locations, (c) five sowing times, and (d) historic and present climate. Average deviation of maximum temperature from the *y = x* line for (f) 11 locations, (g) five sowing times, and (h) historic and present climate. Average deviation of minimum temperature from the *y = x* line for (j) 11 locations, (k) five sowing times, and (l) historic and present climate. In bcd, fgh, jkl error bars are two standard errors and p from ANOVA. In bjf, locations are from north to south.

Deviations of the water stress index from the *y = x* line varied with location, sowing time, and climate (Fig. 7bcd). Owing to the large data set, all three sources of variation returned p < 0.0001, but the size of the effect ranked location > sowing time > climate (Fig. 7bcd). The deviation was small in Emerald and Hermitage, and largest in Roma due to shallow soil (0.7 m) in this location. The deviations of water stress index from the *y = x* line were larger in more stressful locations (Supplementary Figure 2), indicating a larger adaptive value of drought effect on phenology under more severe drought. Deviations in water stress index from the *y = x* line declined from early to late sown crops (Fig. 7c) and were slightly larger for present compared to historic climate (Fig. 7d). For maximum (Fig. 7fgh) and minimum temperature (Fig. 7jkl), deviations were larger for location than for time of sowing, with little or no variation with climate. The deviations for maximum temperature increased southwards (Fig. 7f) with no latitudinal trend for minimum temperature (Fig. 7j).

## Discussion

In this study we define a new trait, drought effect on phenology, and demonstrate it is genotype-dependent, with a broad-sense heritability of 0.61 for our combination of genotypes and environments (Fig. 4); it comprises temperature-dependent and temperature-independent components associated with distinct genomic regions (Fig. 5) that map to genes related to floral development, hormone signalling and abiotic stress signalling (Supplementary Table 4, Table 2), and is involved in a site-dependent adaptive trade-off whereby a hastening of reproductive development with water deficit reduces the risk of drought and heat stress during the critical period of yield formation at the expense of cold risk (Fig. 6, 7, Supplementary Fig. 2).

### Candidate genes involved in the development of reproductive organs, hormone signalling and abiotic stress signalling were under selection for drought effect on phenology

F_ST_ genome scan reliably identifies genomic regions associated with ecologically and agronomically important traits in small populations (Barr *et al*., 2021; Jordan *et al*., 2015; Li *et al*., 2017; Sadras *et al*., 2016; Van Bocxlaer, 2017; Xu *et al*., 2012). Genomic regions associated with *DEP* in early- or late-sown crops, but not in both, were assumed to reflect the temperature-dependent component of the trait. There were 232 predicted genes located within 250kb of the peaks for *DEP* in early sowing, including genes involved in floral development, hormone signalling and abiotic stress signalling. For example, the embryonic flower 1 (*EMF1*) gene is a repressor of the floral meristem determinacy gene AGAMOUS during vegetative development in *Arabidopsis thaliana* via polycomb group (PcG)–mediated gene silencing (Aubert *et al*., 2001; Calonje *et al*., 2008). Another gene, BRANCHED1 (*BRC1/TCP18*), interacts with the florigen proteins FLOWERING LOCUS T (FT) to repress the floral transition of the axillary meristems in *A. thaliana* (Niwa *et al*., 2013). There were 240 predicted genes located within 250kb of the F_ST_ peaks for *DEP* in late sowing (Supplementary Table 4). Some of them are invovled in floral development and abiotic stress signalling. Among them, EMBRYONIC FLOWER 2 (*EMF2*) has been revealed to encode a novel zinc finger protein that repress reproductive development by changing flowering time and shoot morphogenesis (Yoshida *et al*., 2001). Further, EMF2 protein was recruited to interact with several key regulatory genes (*ABI3, LOV1*, and *FLC*) involved in FLC-mediated flowering pathway, seed development, and cold signalling (Kim *et al*., 2010). We also identified the *NPGR1* gene (No Pollen Germination1 Related), one of the three closely related calmodulin-binding proteins, which is essential for pollen germination (Golovkin and Reddy, 2003). Additionally, there are four ethylene-responsive transcription factors: *ERF110, ERF1A, ERF034*, and *WIN1*. They all contain a DNA binding domain (AP2 domain) that could bind to genes that respond to abiotic and biotic stresses (Muller and Munne-Bosch, 2015). Some members of the ERF family regulate floral development through environmental stimuli or hormones (Krizek, 2009; Licausi *et al*., 2013). One key, well-characterised member, *ERF1*, is involved in ethylene signalling (Muller and Munne-Bosch, 2015), and drought and heat stress (Cheng *et al*., 2013). Recently, ERF1 was shown to associate with a delay in *Arabidopsis* flowering/floral initiation through direct inhibition of the expression of the key floral integrator *FT* (Chen *et al*., 2021). Of interest, some ERFs play a role in plant-pathogen relations providing further biological links between phenological development and disease resistance. For example, ERF5 and ERF6 are positive regulators of JA-mediated defence and their constitutive expression increased resistance of *A. thaliana* to the fungal necrotroph *Botrytis cinerea* (Moffat *et al*., 2012). *Magnaporthe oryzae*, the causal agent of rice blast disease, strongly induced RiceOsERF922, encoding an APETELA2/ethylene response factor (AP2/ERF) (Liu *et al*., 2012). The phenotypic and genetic links between phenological development and disease tolerance have been and remain critical for chickpea adaptation, as discussed in the next section.

F_ST_ peaks on chromosome 8 common to early and late sowing are nearby (∼100kb) with high linkage disequilibrium. The 250kb regions surrounding the two peaks overlapped and contain genes associated with a temperature-independent drought effect on phenology. In this region, a cluster of four genes was identified that relates to floral and reproductive development of *Arabidopsis*. One of them is the HAESA-LIKE2 (*HSL2*) receptor-like kinase that activates floral organ abscission, a cell-separation process that allows plants to develop their organ shape in response to developmental cues and environmental stress (Gubert and Liljegren, 2014). The other two genes, *BAM1* and *BAM2*, play important roles in ovule and pollen formation (DeYoung and Innes, 2006; Hord *et al*., 2006). Their functions appear to be overlapping and opposite to that of CLAVATA *1*, a key protein kinase in regulating the development of shoot and flower meristems. The *BAM2* gene in the chickpea reference genome Kabuli v2.6.3 is incomplete, with only 200bp of the total ∼3000bp length. It is unclear whether this is an assembly error or a loss of function deletion that often arises from gene duplication. We also identified the AP2 transcription factor AINTEGUMENTA (*ANT*) gene, which is involved in regulating ovule and female gametophyte development and promotes early floral primordia growth through stimulating cell growth in floral organs (Elliott *et al*., 1996; Krizek, 2009; Krizek *et al*., 2021). Interestingly, the upstream regulator of ANT, BRASSINAZOLE-RESISTANT 1 (*BZR1*), locates in another F_ST_ peak for *DEP* in early sowing (1.5Mb away from each other). The *BZR1* mutant can increase the number of ovules and seeds in *Arabidopsis* through the brassinosteroid signalling pathway (Huang *et al*., 2013). Also, *BZR1* was able to up-regulate the expression of ANT by binding to its promoter sequence and is thus involved in plant reproductive development (Huang *et al*., 2013).

We did not find any genomic region under selection for *DEP* overlapping the typical genes reported for flowering time in response to photoperiod and temperature in chickpea (Gaur *et al*., 2015; Gursky *et al*., 2018). The corollary of this putative genetic independence is that phenotypes combining slow or fast development, as mediated by photoperiod and temperature, and small or large responsiveness to drought can be tailored to target environments.

### Drought effect on phenology may be adaptive and involves trade-offs

To interpret the assumed adaptive value of drought effect on phenology, we first outline the interplay between phenology, climate, and Ascochyta blight, as drivers of evolution, and early and contemporary cultivation of chickpea (Abbo *et al*., 2002; Abbo *et al*., 2003; Abbo *et al*., 2008; Daba *et al*., 2016a; Kumar and Abbo, 2001; Li *et al*., 2017; Lichtenzveig *et al*., 2006). In the Near East’s archaeological record, chickpea first appears with the “large-seeded legumes” about 13,000 Cal BP, followed by a gap of about 3,000 years, and its re-appearance in the Bronze age. The gap has been attributed to Ascochyta blight, which devastated autumn-sown crops, and the re-appearance of the crop associated with the shift from autumn to spring sowing to avoid disease (Abbo *et al*., 2003). The selective pressure in favour of a spring-summer phenotype has reduced or eliminated vernalization requirements in cultivated chickpea in comparison to both wild *Cicer spp* and the companion foundational crops in the Levante (Abbo *et al*., 2002; Abbo *et al*., 2003; Berger *et al*., 2005; Kumar and Abbo, 2001; Pinhasi Van-Oss *et al*., 2016). In western Canada where short growing season and Ascochyta blight challenge contemporary chickpea production, a collection of recombinant inbred lines showed negative correlations between days to flowering and Ascochyta blight resistance, and revealed clusters of QTL for days to flowering and blight resistance that partially overlap on chromosomes 3 (8.6–23.11 cM) and 8 (53.88–62.33 cM) (Daba *et al*., 2016a). In Australia where the current crop is autumn-sown, the legacy of selection for summer growth habit is apparent in two traits: slow canopy growth under low temperature, compared to other autumn-sown pulses such as field pea, and a high rate of pod abortion in cool springs (Berger *et al*., 2005; Lake *et al*., 2016). These traits determine that severe drought is unlikely before flowering (Chauhan *et al*., 2017; Lake *et al*., 2016).

Against this ecological and agronomic background, we modelled the relative risk of drought, heat, and cold stress in the critical period for two contrasting genotypes. We found that drought effect on phenology consistently alleviates the severity of drought during the critical period of yield formation, and that the adaptive value of this trait is higher in more stressful environments, e.g., shallow soil and late sowing. The consistent alleviation of drought stress with hastened flowering relates to the pattern of drought in these environments and cannot be extrapolated to other conditions such as intermittent drought (Jordan and Miller, 1980; Schwinning and Ehleringer, 2001; Tardieu, 2012). Owing to the strong seasonality of rainfall in eastern Australia, crops mostly rely on in-season rainfall in the southern locations of our transect, and rely primarily on stored soil water in the northern locations (Sadras and Rodriguez, 2007); this geographical divide mirrors the early selective pressures for autumn- and spring-sown crops outlined above. In both cases, slow-growing crops and high availability of soil water (north) or winter rainfall (south) combine to return typically unstressed conditions early in the season, and a characteristic terminal drought with declining water availability during critical reproductive stages in spring as illustrated in Fig. 1a (Chauhan *et al*., 2017; Lake *et al*., 2016). The reliability of environmental cues influences the evolution of signalling pathways in plants (Aphalo and Sadras, 2022); a predictable drought pattern over long time scales (millennia) is thus consistent with the selection for hastened flowering in response to drought. The responsive genotype had lower maximum temperature during the critical period, that can further contribute to seed yield, and lower minimum temperature, which could compromise pod set (Berger *et al*., 2012; Devasirvatham *et al*., 2012).

## Conclusion: biological, agronomic, and modelling implications, and further research

The combination of experiments in realistic field conditions, F_ST_ genome scan, and modelling, highlighted the agronomic importance of water deficit effect on chickpea phenology, an overlooked trait. Controlled experiments combining water and thermal regimes, ideally with isogenic lines, are needed to unequivocally assess the agronomic value, i.e., seed yield response, of drought effect on phenology. The genetic variation in our small sample and the relatively high heritability of this trait suggest further research is warranted for breeding applications. Several candidate genes homologous to *Arabidopsis thaliana* involved in the development of floral and reproductive tissues and stress responses were identified with a putative role in the response of chickpea phenology to drought. Of note, the core genetic regulatory network canalizing the flowering signals to the decision to flower in *A. thaliana* partially holds in chickpea (Gursky *et al*., 2018). Further molecular function experiments are needed to establish their roles for this trait. Irrespective of the actual genes and metabolic pathways, we demonstrated a strong influence of drought on reproductive development that needs to be incorporated in both genetic models (Gursky *et al*., 2018) and phenotype models that simulate crop phenology, carbon and water dynamics, growth and yield (Chauhan *et al*., 2019). Models exclusively based on temperature and photoperiod are bound to return biased predictions of phenology, and hence unreliable predictions of crop adaptation to future, drier climates in Australia and elsewhere.

## Acknowledgements

We thank the Grains Research and Development Corporation for funding (projects UOT1909-002RTX, DAS00140), and Francisco Sadras for expert contribution to data analysis.

## Author Contributions

YL: carried out F_ST_ analysis, wrote part of the paper; YSC: modelled phenology and water stress; LL: carried out field experiment, contributed data analysis; JT: carried out statistical analysis; VOS: developed the concept, designed the experiment, analysed data, wrote the manuscript.

## Supplementary data

Supplementary data are available at JXB online.

**Supplementary Table 1.** Genetic parameters to model phenological development of chickpea. Parameters are based on measurements and calibrations with PBA Boundary, a locally adapted, widely used cultivar in commercial crops.

| Parameter | Description | Unit | Value or Range |
| --- | --- | --- | --- |
| tt_emerg_to_endjuv | TT from emergence to end of juvenile phase | °Cd | 650 |
| cum_vernal_days | cumulative vernal days | d | 0 to 100 |
| est_days_emerg_to_init | Days from emergence to floral initiation | d | 83 |
| x_pp_endjuv_to_init | Photoperiod | h | 10.7 to 17.0 |
| y_tt_endjuv_to_init | TT from end juvenile to floral initiation | °Cd | 446 |
| x_pp_init_to_flower | Photoperiod | h | 1 to 24 |
| y_tt_init_to_flower | TT from initiation to flowering | °Cd | 33 |
| x_pp_flower_to_start_grain | Photoperiod | h | 1 to 24 |
| y_tt_flower_to_start_grain | TT from flowering to start grain fill | °Cd | 450 |
| x_pp_start_to_end_grain | Photoperiod | h | 1 to 24 |
| y_tt_start_to_end_grain | TT from start grain fill to end grain fill | °Cd | 690 |
| tt_end_grain_to_maturity | TT from end grain fill to maturity | °Cd | 60 |

**Supplementary Table 2.** Analysis of variance of thermal time from sowing to flowering, carbon isotope composition δ^13^C, and seed weight.

| Source of variation | Df | Thermal time to |  |  |  |  |  |
| --- | --- | --- | --- | --- | --- | --- | --- |
| | | flowering | | $\delta^{13}\text{C}$ | | Seed weight | |
|  |  | Wald<br>statistic | Pr(Chisq) | Wald<br>statistic | Pr(Chisq) | Wald<br>statistic | Pr(Chisq) |
| Intercept | 1 | 577198.6 | < 2.2E-16 | 333960.9 | < 2.2E-16 | 109674.2 | < 2.2E-16 |
| Season | 1 | 1176.3 | < 2.2E-16 | 76.0 | < 2.2E-16 | 0.6 | 0.43 |
| Sowing time | 1 | 6684.2 | < 2.2E-16 | 459.5 | < 2.2E-16 | 24.1 | 9.08E-07 |
| Water regime | 1 | 228.3 | < 2.2E-16 | 47.6 | 5.19E-12 | 25.4 | 4.70E-07 |
| Genotype | 19 | 3354.7 | < 2.2E-16 | 115.7 | 6.66E-16 | 20406.5 | < 2.2E-16 |
| Season:Sowing time | 1 | 89.8 | < 2.2E-16 | 0.0 | 0.92 | 2.1 | 0.15 |
| Season:Water regime | 1 | 1.7 | 0.19 | 4.2 | 0.04 | 9.3 | 0.002348 |
| Sowing time:Water regime | 1 | 84.3 | < 2.2E-16 | 3.5 | 0.06 | 4.8 | 0.028273 |
| Season:Genotype | 19 | 106.4 | 3.71E-14 | 17.1 | 0.58 | 140.5 | < 2.2E-16 |
| Sowing time:Genotype | 19 | 137.6 | < 2.2E-16 | 15.0 | 0.72 | 30.2 | 0.049 |
| Water regime:Genotype | 19 | 165.2 | < 2.2E-16 | 44.0 | 0.00094 | 77.5 | 4.93E-09 |
| Season:Sowing time:Water regime | 1 | 6.7 | 0.0099 | 0.7 | 0.41 | 2.5 | 0.11 |
| Season:Sowing time:Genotype | 19 | 77.5 | 4.93E-09 | 25.4 | 0.15 | 58.1 | 7.74E-06 |
| Season:Water regime:Genotype | 19 | 48.2 | 0.00024 | 20.5 | 0.37 | 59.8 | 4.12E-06 |
| Sowing time:Water regime:Genotype | 19 | 104.9 | 6.91E-14 | 25.5 | 0.14 | 19.8 | 0.41 |
| Season:Sowing time:Water regime:Genotype | 19 | 24.4 | 0.18 | 22.2 | 0.27 | 24.2 | 0.19 |
| Residual (MS) | NA | NA | NA | NA | NA | NA | NA |

**Supplementary Table 3.** Analysis of variance of drought effect on phenology.

| Source of variation | Df | Wald statistic | Pr(Chisq) |
| --- | --- | --- | --- |
| Intercept | 1 | 386.9 | < 2.2E-16 |
| Season | 1 | 72.3 | < 2.2E-16 |
| Sowing date | 1 | 117.0 | < 2.2E-16 |
| Genotype | 19 | 91.4 | 1.87E-11 |
| Season:Sowing date | 1 | 0.2 | 0.63 |
| Season:Genotype | 19 | 43.7 | 0.0010 |
| Sowing date:Genotype | 19 | 88.0 | 7.50E-11 |
| Season:Sowing date:Genotype | 18 | 28.2 | 0.059 |
| Residual (MS) | NA | NA | NA |

Supplementary Table 4. Genes located within 250kb of the genomic regions under selection (top 0.1% F_ST_) for *DEP*. See attached Excel file.

**Supplementary Table 5.** Summary statistics of frequency distributions of flowering time, water stress index and temperature for responsive and unresponsive genotypes. Histograms are in Figure 6.

| Variable | Parameter | Responsive | Unresponsive |
| --- | --- | --- | --- |
| Time to flowering<br>(d) | Mean | 73 | 89 |
|  | Std. Dev. | 11.5 | 17.5 |
|  | Minimum | 47 | 55 |
|  | Maximum | 110 | 148 |
|  | Coef. Var. | 0.16 | 0.20 |
|  | Skewness | 0.26 | 0.66 |
|  | Kurtosis | -0.18 | -0.02 |
|  | Median | 73 | 87 |
| Thermal time to flowering<br>(°Cd) | Mean | 878 | 1105 |
|  | Std. Dev. | 48.5 | 110.5 |
|  | Minimum | 822 | 846 |
|  | Maximum | 1051 | 1585 |
|  | Coef. Var. | 0.06 | 0.10 |
|  | Skewness | 1.29 | 0.43 |
|  | Kurtosis | 0.43 | 0.04 |
|  | Median | 856 | 1095 |
| Water stress index<br>(unitless) | Mean | 0.978 | 0.927 |
|  | Std. Dev. | 0.065 | 0.127 |
|  | Minimum | 0.377 | 0.293 |
|  | Maximum | 1 | 1 |
|  | Coef. Var. | 0.067 | 0.137 |
|  | Skewness | -4.27 | -2.23 |
|  | Kurtosis | 21.31 | 4.77 |
|  | Median | 1 | 0.993 |
| Maximum temperature<br>(°C) | Mean | 22.3 | 24.2 |
|  | Std. Dev. | 3.2 | 2.9 |
|  | Minimum | 14.2 | 16.1 |
|  | Maximum | 31.7 | 33.5 |
|  | Coef. Var. | 0.14 | 0.12 |
|  | Skewness | 0.03 | 0.17 |
|  | Kurtosis | -0.43 | -0.30 |
|  | Median | 22.3 | 24.1 |
| Minimum temperature<br>(°C) | Mean | 7.1 | 8.7 |
|  | Std. Dev. | 2.4 | 2.4 |
|  | Minimum | -0.1 | 1.5 |
|  | Maximum | 16.0 | 16.6 |
|  | Coef. Var. | 0.34 | 0.28 |
|  | Skewness | 0.22 | 0.16 |
|  | Kurtosis | -0.21 | -0.17 |
|  | Median | 7.0 | 8.7 |
| Mean temperature<br>(°C) | Mean | 14.7 | 16.4 |
|  | Std. Dev. | 2.50 | 2.35 |
|  | Minimum | 8.9 | 10.8 |
|  | Maximum | 22.7 | 24.3 |
|  | Coef. Var. | 0.17 | 0.14 |
|  | Skewness | 0.28 | 0.34 |
|  | Kurtosis | -0.48 | -0.35 |
|  | Median | 14.6 | 16.3 |

**Supplementary Figure 1.**
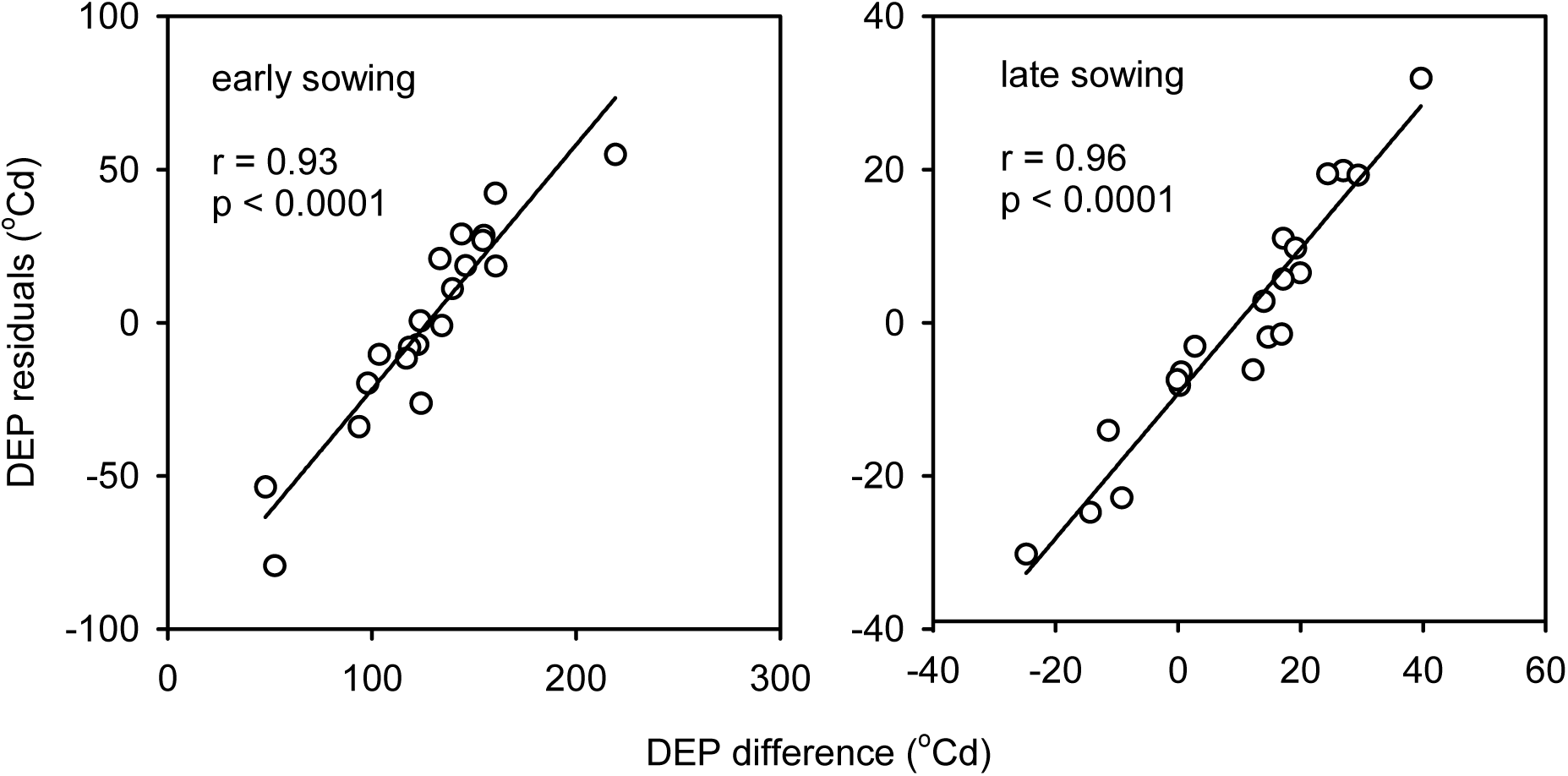
Comparison of drought effect on phenology (*DEP*) calculated with difference- and residual-based approaches.

**Supplementary Figure 2.**
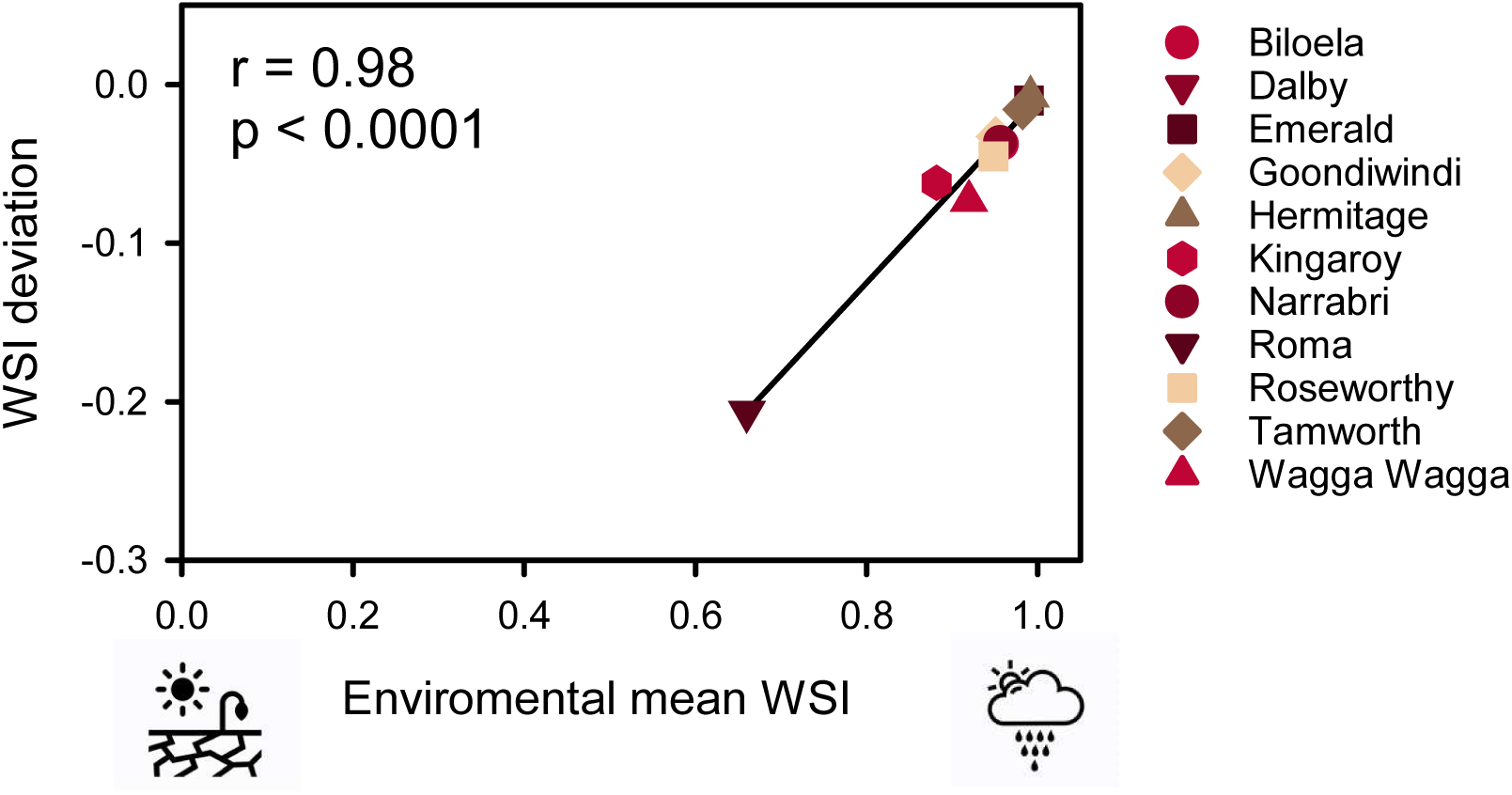
*WSI* deviation from the *y = x* line in Figure 7a as a function of environmental mean *WSI*, representing the average water stress index in the critical period for each location, averaged across other factors (65 years, 5 sowing dates, 2 initial soil waters, 2 genotypes). The *WSI* ranges from 1 (no stress) to 0 (no growth). The line is the least squares regression.

## Data availability

Dryad, Dataset, https://doi.org/10.5061/dryad.tx95x6b0f

## Conflicts of interest

The authors declare no conflicts of interest.

## Footnotes

1 https://www.longpaddock.qld.gov.au/silo/point-data/

## Notes

### Competing Interest Statement

The authors have declared no competing interest.

